# Gut microbial features can predict host phenotype response to protein deficiency

**DOI:** 10.1101/398248

**Authors:** Guadalupe Navarro, Anukriti Sharma, Lara R. Dugas, Terrence Forrester, Jack A. Gilbert, Brian T. Layden

## Abstract

Malnutrition remains a major health problem in low and middle income countries. During low protein intake, < 0.67 g/kg/day, there is a loss of nitrogen (N_2_) balance, due to the unavailability of amino acid for metabolism and unbalanced protein catabolism results. However, there are individuals, who consume the same low protein intake, and preserve N_2_ balance for unknown reasons. A novel factor, the gut microbiota, may account for these N_2_ balance differences. To investigate this, we correlated gut microbial profiles with the growth of four murine strains (C57Bl6/J, CD-1, FVB, and NIH-Swiss) on protein deficient (PD) diet. Results show that a PD diet exerts a strain-dependent impact on growth and N_2_ balance as determined through analysis of urinary urea, ammonia and creatinine excretion. Bacterial alpha diversity was significantly (p < 0.05, FDR) lower across all strains on a PD diet compared to normal chow (NC). Multi-group analyses of the composition of microbiomes (ANCOM) revealed significantly differential microbial signatures between the four strains independent of diet. However, mice on a PD diet demonstrated differential enrichment of bacterial genera including, *Allobaculum* (C57Bl6/J), *Parabacteroides* (CD-1), *Turicibacter* (FVB), and *Mucispirillum* (NIH-Swiss) relative to NC. Additionally, statistical model fitting revealed that the relative abundance of genera such as *Bifidobacterium, Ruminococcus,* and *Lactobacillus* were significantly positively correlated with body weight, while *Anaerofustis, Roseburia*, and *Bilophila* were significantly positively correlated with ammonia excretion. Taken together, these results suggest a potential relationship between the specific gut microbiota, N_2_ balance and animal response to malnutrition.

## Introduction

Both severe chronic malnutrition (−3 standard deviations (SD) below median World Health Organization (WHO) height-for-age score) and severe acute malnutrition (−3 SD below WHO weight-for-height score) annually affect approximately 200 million children worldwide, with almost half of the deaths in children under five years being directly or indirectly attributable to insufficient nutrition (3). Severe childhood malnutrition can be lethal, with mortality rates between 10 and 50% depending on the setting and the appropriateness of clinical care available (9). In the longer term, both severe acute malnutrition (wasting with and without edema) and severe chronic malnutrition, also known as stunting, can impair health, increasing the likelihood of obesity and cardiometabolic disease as well as neurocognitive impairment throughout the life cycle (12,13,24).

The determinants of malnutrition include both macro- and micronutrient deficiencies often accompanied by bacterial, viral and parasitic infectious diseases. Macronutrient deficiency, especially when derived from inadequate intakes of dietary protein to meet metabolic demand, is thus a major contributor to stunting and wasting. Surprisingly, these are not only common in low and middle-income countries, but also affect sub-populations in more developed countries (27). Starting from any plane of nutrition, a reduction in dietary protein intake results in inadequate supply of amino acids with respect to initial metabolic demand. The response of whole body protein metabolism to this unmet demand for amino acids will be unbalanced catabolism of lean body mass. Whether or not balance is achieved during protein restriction depends on several variables, including dietary protein quality, metabolic demand of the host for amino acids, as well as individual variation in the adaptive capacity for maintaining N_2_ balance. During dietary protein restriction, although protein catabolism releases amino acids into the whole body amino acid pool to support integrated metabolism, there is nonetheless a mismatch of amino acid requirements due to the restricted supply. Consequently, amino acids oxidation rises with metabolism of the N_2_ moiety to urea and ammonia in the liver and kidney which are excreted in the urine. A proportion of the urea enters the gastrointestinal tract where it is made available for microbial protein and amino acid metabolism. There is evidence to support the transfer of bacterially manufactured amino acids synthesized by gut bacteria to the host for use in protein metabolism (2,23,28).

Determining the microbial contribution to whole-body protein requirement may have significant implications for understanding the variation in human adaptation to reduced dietary protein intake. We posit that one of the key components of this inherent adaptive capacity to low protein diet is the ability of the gut microbiota to provide amino acids to the host. In this study, we characterized how gut microbial profiles correlate with murine model growth on a protein-restricted diet. Specifically, we leveraged the diverse microbial communities found in 4 different murine models to determine how the microbiota correlate with maintenance of positive N_2_ balance during marginal protein intake. We hypothesized that the differential microbiota associated with each mouse strain would correlate with the differential growth and N_2_ balance on a protein restricted diet.

## Materials and Methods

### Experimental animals

In this study, four strains of male mice (C57Bl6/J, CD-1, FVB, and NIH-Swiss, n=10 for each strain) were obtained at 10 weeks of age, individually housed, allowed to acclimate to facility for 1 week. Following this, the mice were placed either on normal chow (NC, 20% protein) or protein deficient diet (PD, 6% TD.90016; Envigo) and followed for 4 weeks. Weekly fecal samples and weights were captured in all groups. Following 4 weeks of PD or NC, urine was collected for assessment of N_2_ balance, and the mice were euthanized. Plasma, serum, liver, and fecal material were collected. The study was approved by Jesse Brown VA Medical Center IACUC in accordance with the NIH Guide for the Care and Use of Animals.

### Nitrogen balance measurements

Urine and serum samples collected were stored at − 80°C. Ammonia was measured using a commercially available kit (AA0100; Sigma). Urine creatinine was assessed using a colorimetric assay kit (ab65340; Abcam). Urine and serum urea were measured using a urea assay kit (MAK006; Sigma).

### 16S rRNA gene sequencing

Bacterial DNA was extracted using the PowerSoil DNA isolation kit (MoBio Laboratories) following the protocol of Flores (11). For the microbiota analysis, bacterial DNA was extracted from the fecal samples (0.25 g). The V4 region of the 16S rRNA gene (515 F-806R) was amplified with region-specific primers that included the Illumina flowcell adapter sequences and a 12-base barcode sequence. Each 25 µl PCR reaction contained the following: 12 µl of MoBio PCR Water (Certified DNA-Free; MoBio), 10 µl of 5 Prime HotMasterMix (1×), 1 µl of forward primer (5 µM concentration, 200 pM final), 1 µl of Golay Barcode Tagged Reverse Primer (5 µM concentration, 200 pM final), and 1 µl of template DNA (25). The conditions for PCR were as follows: 94°C for 3 min to denature the DNA, with 35 cycles at 94°C for 45 s, 50°C for 60 s, and 72°C for 90 s, with a final extension of 10 min at 72°C. Amplicons were quantified using PicoGreen (Invitrogen) and a plate reader followed by clean up using UltraClean® PCR Clean-Up Kit (MoBio), and then quantification using Qubit (Invitrogen). Samples were sequenced on the Illumina MiSeq platform at the Argonne National Laboratory core sequencing facility according to EMP standard protocols (http://www.earthmicrobiome.org/emp-standard-protocols/its/);

### 16S data analyses

For 16S rRNA analysis, the 16 million paired-end reads generated were joined using join_paired_ends.py script followed by quality-filtering and demultiplexing using split_libraries_fastq.py script in QIIME 1.9.1 (5). Parameters for quality filtering included 75% consecutive high-quality base calls, a maximum of three low-quality consecutive base calls, zero ambiguous bases, and minimum Phred quality score of 3 as suggested previously (4). The final set of demultiplexed sequences were then selected for identifying exact sequence variants (ESVs) using DeBlur pipeline (1). In the pipeline, *de novo* chimeras were analyzed and remove, artifacts (i.e. PhiX) were removed, and ESVs with under 10 reads were removed. The final biom file was also rarefied to 7,565 reads (minimum number of reads) per sample to avoid sequencing bias, which was then analyzed in phyloseq package in R (19). Random forest supervised learning models were used to estimate the predictive power of microbial community profiles for diet i.e. NC or PD. The supervised learning was performed employing Out-Of-Bag (OOB) sample sets in RandomForest package in R. Training was accomplished in RandomForest with bootstrapping at 1000 trees and prediction accuracy (1-OOB) was estimated. We also predicted and annotated the most important bacterial ESVs for differentiating between mice on PD or NC diet using RandomForest in R.

### Statistical analysis

The data for body weight, ammonia and creatinine excretion were analyzed by Student’s t test and expressed as mean ± standard error of the mean (SEM). Significance threshold was established as p ≤ 0.05. Principal coordinate analysis (PCoA) plots were utilized to analyze all microbiota samples. Microbial alpha diversity (based on Shannon and Inverse Simpson indices) between groupings such as strains and diet type were assessed for significance using a nonparametric two-group t-test over 999 Monte Carlo permutations (6). Beta-diversity was determined using both weighted and unweighted UniFrac distance matrices (17). An unweighted UniFrac distance uses only species presence and absence information and counts the fraction of branch length unique to either community. On the other hand, a weighted UniFrac distance uses species abundance information and weights the branch length with abundance difference. Therefore, while the Unweighted UniFrac distance is most efficient in detecting abundance change in rare lineages, the weighted UniFrac distance can efficiently detect changes in abundant lineages. The differences in alpha and beta diversity indices were then tested for significance using permutational multivariate analysis of variance (PERMANOVA). Analysis of composition of microbes (ANCOM) was used to identify differentially abundant bacterial ESVs between different groups i.e. four strains and different diets (NC vs. PD diet) at a p-value cut-off of 0.05 and Benjamini-Hochberg FDR correction (18). Weighted correlation network analysis (WGCNA) package in R was used to identify clusters (modules) of significantly correlated ESVs (16). To minimize spurious associations during module identification, we transformed the adjacency into Topological Overlap Matrix (TOM) and calculate the corresponding dissimilarity (15). ESVs were organized into modules, using this topological overlap measure as a robust measure of interconnectedness in a hierarchical cluster analysis implemented in WGCNA package in R. To further build associations between the modules and body weight, ammonia and creatinine production, we used eigengene network methodology to identity potential significant associations (p < 0.05). The generalized linear models (GLMs) were run in order to determine potential correlations between selected genera (significantly enriched in each strain type PD group from ANCOM) and body weight, creatinine and ammonia excretion. GLMs were implemented in glm package (in R) using the counts data for the genera using Poisson regression and “log” link. The significance was denoted by each correlation for the clinical parameters was evaluated using ANOVA and “chisq” test to compare nested models. For the GLMs, standardized beta coefficients were plotted (14) to overcome the bias introduced due to varying scales (units) for the three input variables i.e. body weight, creatinine and ammonia excretion.

## Results

### Strain-specific weight and nitrogen balance response to low protein diet

The murine cohorts (n=10 per group) were allocated to either a PD diet or continued on NC diet. Weekly mice body weights of both diet cohorts over 4 weeks for each strain was quantified (**Fig. 1**). Both C57BI6/J and NIH-Swiss mice failed to maintain weight gain on PD compared to their age matched controls on NC. Specifically, PD body weight for both C57Bl6/J mice and NIH-Swiss mice was significantly less during week 3 and 4 relative to their matched NC mice. CD-1 and FVB mice show comparable weight gain between the PD and NC groups (**Fig 1 A-D**).

**Figure 1:**
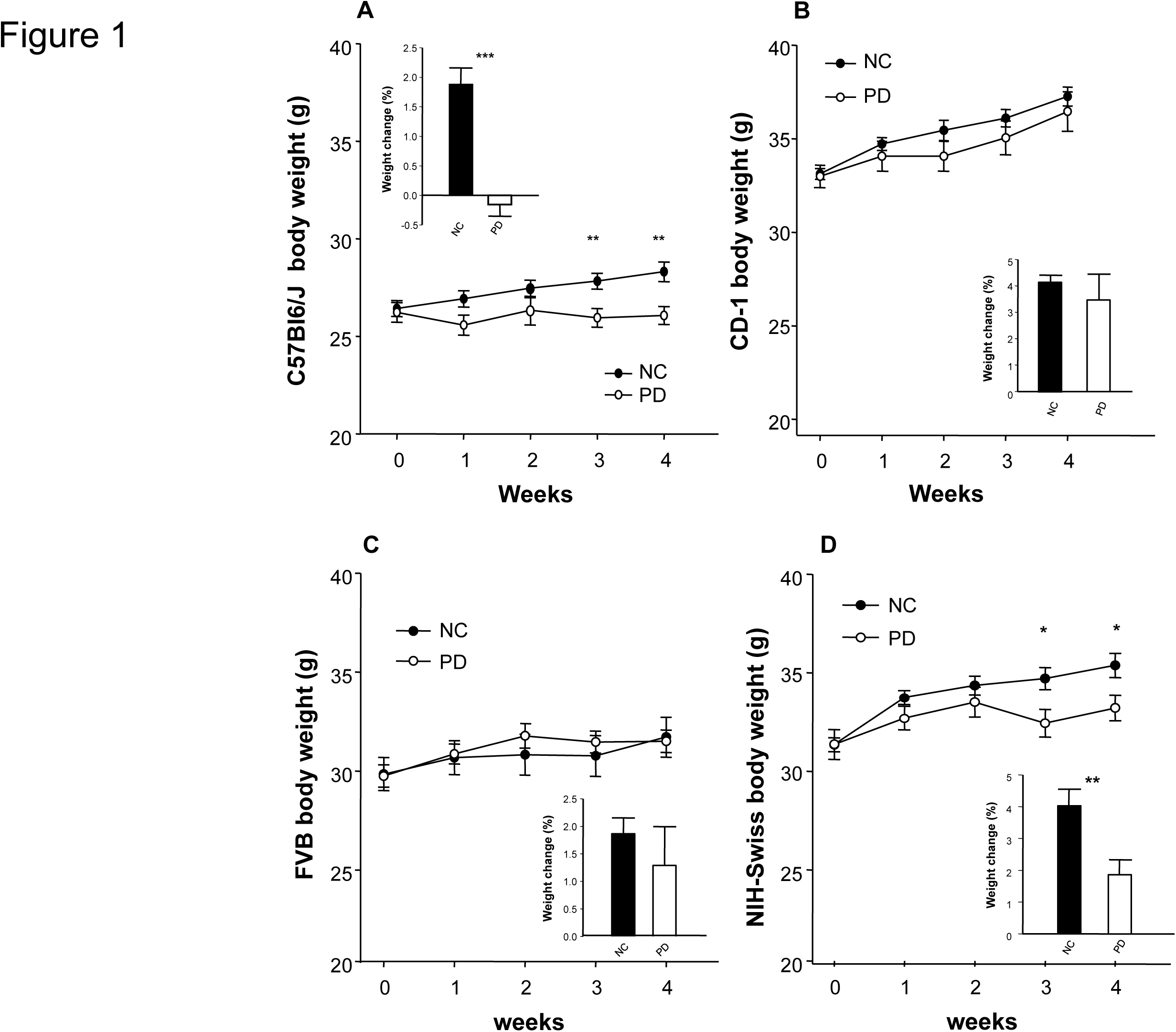
Effects of a low protein diet (PD) on weekly body weights compared to normal chow (NC) diet C57BI6/J (A), CD-1 (B), FVB (C), and NIH-Swiss mice (D) during 4 weeks protein deficient (PD) diet (n=10). Values represent the mean ± sem *p < 0.05, **p < 0. 0.01, one-way ANOVA.

At 4 weeks of diets, urea, ammonia and creatinine were measured in urine as markers of N_2_ balance in the 4 murine strains. The PD diet was associated with significantly diminished urea concentration in urine and serum for both CD-1 and C57Bl6/J relative to NC mice (**Fig 2 A-B**), while in NIH-Swiss mice, urea was only significantly reduced in serum on PD compared to NC (**Fig 2 A-B**). Urinary ammonia concentration was significantly reduced on the PD diet compared to NC in CD-1 and NIH-Swiss mice (**Fig. 2C**). FVB mice retained similar urea and ammonia concentrations both in urine and serum independent of diet (**Fig 2 A-D**); similarly, C57Bl6/J mice showed no significant reduction in urine and serum ammonia on PD compared to NC (**Fig 2C-D**). In fact (**Fig 2D**), serum ammonia levels, while lower on a PD, were not significantly reduced compared to NC for C57BI6/J, CD-1, and NIH-Swiss mice. While the same is true for FVB mice, it is interesting to note that serum ammonia concentration was slightly greater on the PD compared with NC (**Fig 2D**). Therefore, the PD diet had a murine strain-dependent impact on urea excretion for both urine and serum, while urinary ammonia was only significantly different for some strains, and not serum. And finally, renal creatinine excretion was quantified in urine at a single time point. The PD diet was associated with a consistent, but non-significant decrease in creatinine on a PD diet compared to the NC diet for all 4 mice strains (**Fig 2E**).

**Figure 2:**
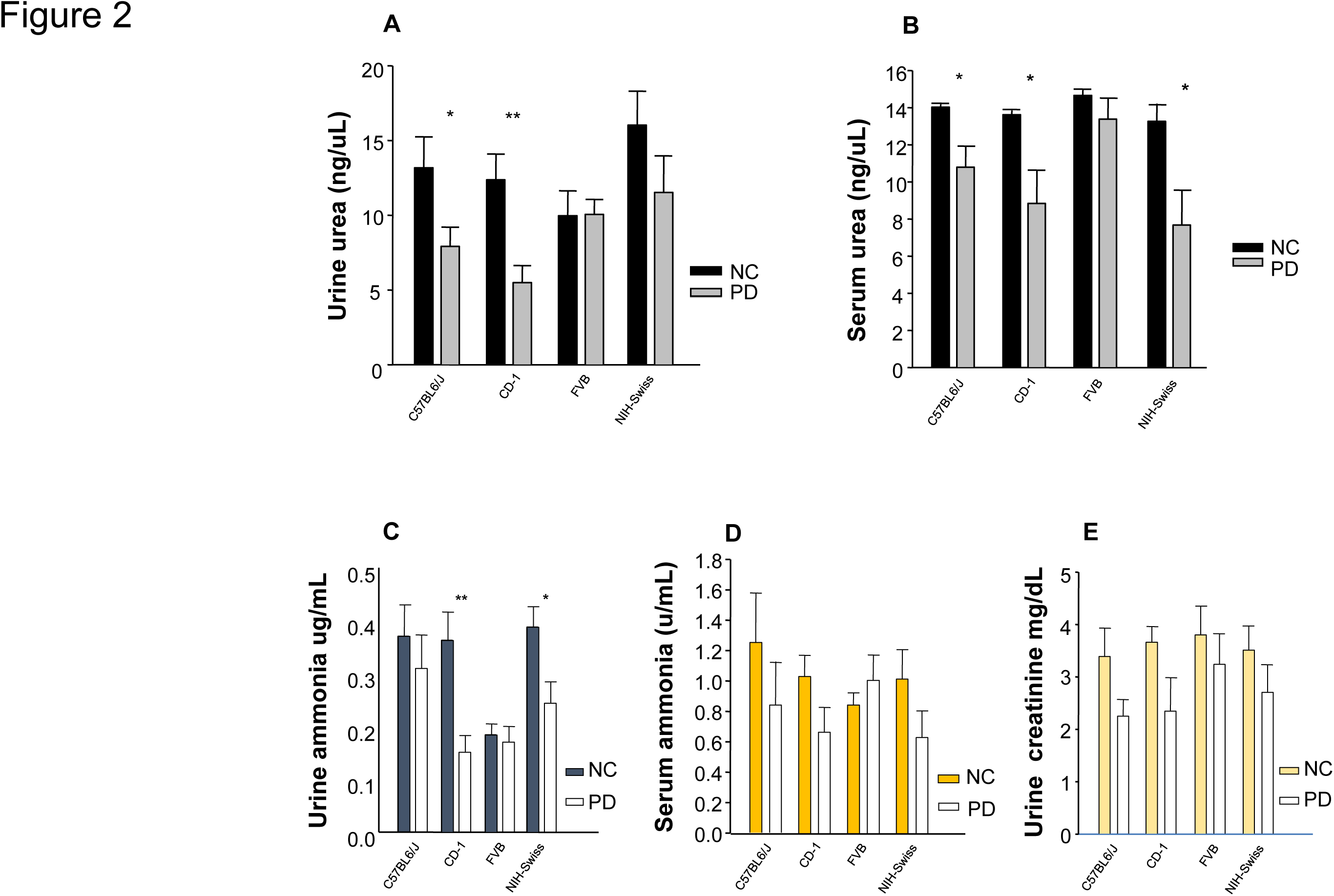
In each strain of mice, the nitrogen response is shown, including urine urea concentration (A), serum urea concentration (B), urinary ammonia (C), serum ammonia (D), and urinary creatinine (E), while the mice were on either 6 % protein diet (PD), as compared to the NC diet. N=10, 15-weeks old mice. Values represent the mean ± sem *p < 0.05, **p < 0. 0.01, one-way ANOVA.

### Gut microbial diversity and composition associate with low protein diet

Bacterial alpha diversity (determined using 16S rRNA amplicon sequencing) was significantly lower (p_FDR_ < 0.05) on a PD diet compared to a NC diet for all 4 murine strains, albeit for different metrics (**Fig. 3A**). Beta-diversity was also significantly different between the strain types and diets using both Weighted and Unweighted UniFrac distance (PERMANOVA; p<0.05). However, using PCoA plots the groups were only visibly differentiated (i.e. group specific clustering) with an unweighted UniFrac distance metric, suggesting that rare bacterial taxa contribute more toward observed differences. In the Unweighted UniFrac PCoA plot, the FVB and NIH-Swiss samples clustered together by diet, i.e. the FVB, NIH-Swiss samples on PD diet clustered closely together and the FVB, NIH-Swiss samples on NC diet clustered closely (**Fig. 3B**). Similarly, C57BI6/J and CD-1 clustered together separately also by diet (**Fig. 3B**). Therefore, while C57BI6/J and CD1 are more similar to each other than FVB and NIH-Swiss, diet is still a key driver of differences in the abundance and composition of less abundant bacterial strains.

**Figure 3:**
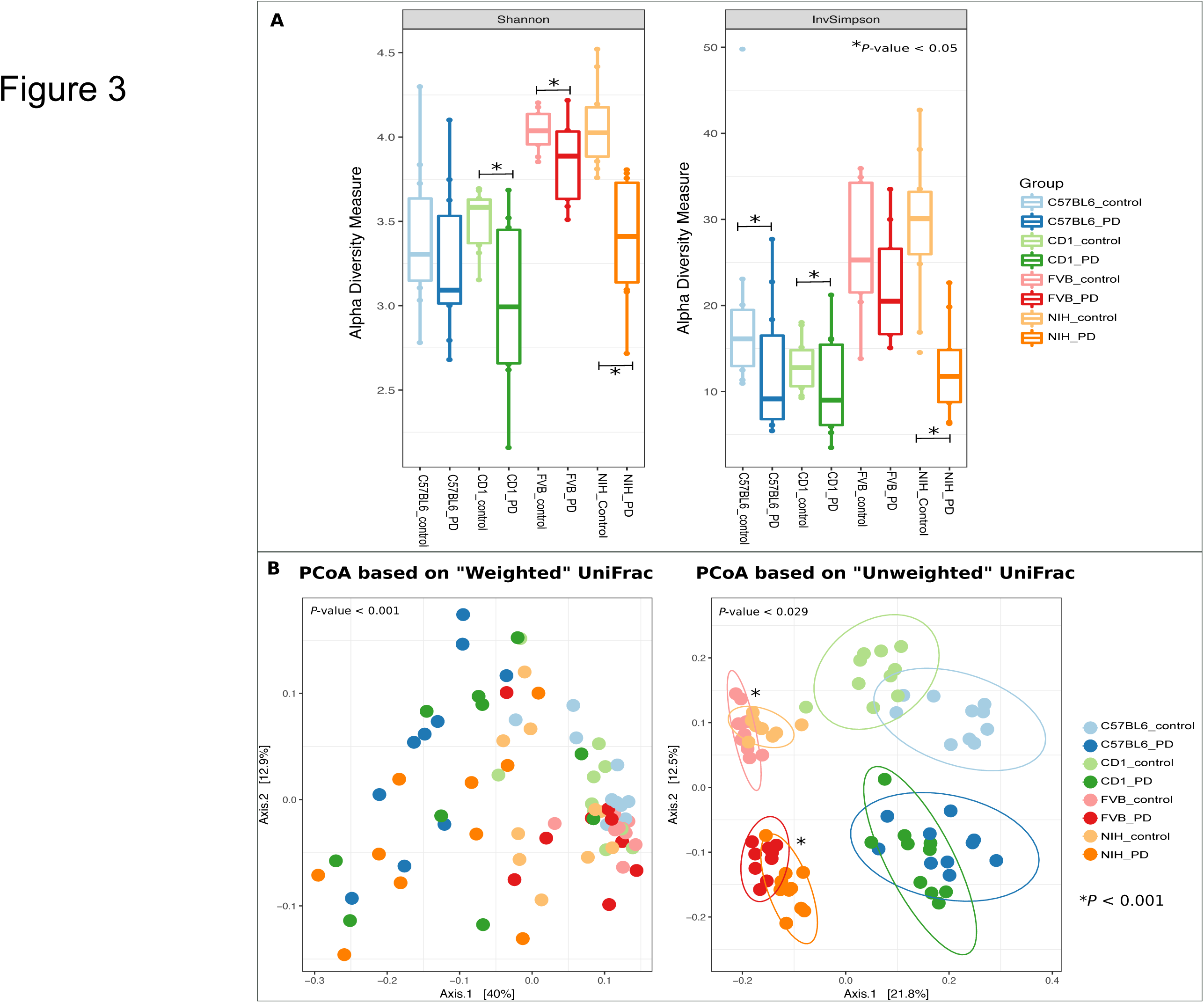
Alpha and beta diversity comparisons between the control and PD mice of the 4 strains i.e. C57BI6/J, CD-1, NIH-Swiss, and FVB. Shannon and Inverse Simpson metrics were used to compare control and PD mice across the 4 strains (A). PERMANOVA test was used to test significance of the variation observed. The asterisks show significant difference between the 2 groups. The PCoA shows the clustering pattern between the control and PD mice based on both weighted and unweighted UniFrac metrics (B). The unweighted distance based PCoA revealed distinct clustering pattern between different groups.

Using a multi-group ANCOM analyses, the bacterial genera that significantly differed (p_FDR_ < 0.05) in proportion between the 4 mice strains on NC diet, were identified (**Table S1**). C57BI6/J mice on NC diet were enriched (p_FDR_ < 0.05) for ESVs belonging to genera *Sutterella* and *Turicibacter*, FVB mice were enriched (p_FDR_ < 0.05) for ESVs belonging to order Bacteroidales, family Rickenellaceae, and genus *Odoribacter* (**Table S1**). Similarly, CD-1 mice on NC diet were significantly (p_FDR_ < 0.05) enriched for ESVs belonging to genus *Bacteroides*, while NIH-Swiss mice were significantly (p_FDR_ < 0.05) enriched for ESVs belonging to genus *Parabacteroides* (**Table S1**).

Next, the impact of diet type (PD vs. NC) on the relative proportions of specific bacterial genera were determined using ANCOM. In C57BL6/J, ESVs from *Bacteroides, Mucispirillum, Coprococcus, Allobaculum, and Akkermansia* were significantly (p_FDR_ < 0.05) enriched on the PD diet, while ESVs from genera *Turicibacte*r, *Ruminococcus,* order Bacteroidales, and class Mollicutes were significantly enriched on the NC diet (p_FDR_ < 0.05) (**Table S2**). CD-1 mice showed an enrichment (p_FDR_ < 0.05) in ESVs from *Parabacteroides, Allobaculum, and Akkermansia* on the PD diet, and an enrichment in genus *Odoribacter* and one ESV belonging to family Rickenellaceae were enriched on NC diet (p_FDR_ < 0.05) (**Table S3**). In FVB mice, ESVs belonging to *Prevotella*, *Bacteroides, Parabacteroides, Turicibacter*, and *Allobaculum* were significantly enriched on the PD diet, while genera *Odoribacter, Lactobacillus*, ESV from family Rickenellaceae were significantly enriched in FVB mice on NC diet (p_FDR_ < 0.05; **Table S4**). Finally, in NIH-Swiss mice, ESVs belonging to genus *Bacteroides, Parabacteroides*, *Mucispirillum*, and *Allobaculum,* were significantly (p_FDR_ < 0.05) enriched on the PD diet, while *Prevotella, Lactobacillus, and Turicibacter* were significantly enriched on the NC diet (p_FDR_ < 0.05) (**Table S5**). Therefore, the microbiota of C57Bl6/J and CD1 mice were again found to be more similar, as compared to the FVB and NIH-Swiss which had greater microbial similarity (**Fig. 3B**). Despite these similarities, the bacterial genera that significantly changed in response to the PD diet were different. When just comparing differences between mouse strains on the PD diet alone, *Prevotella* was significantly enriched in FVB, *Bilophila* was enriched in CD-1, *Coprococcus* and *Coprobacillus* were enriched in C57BI6/J, and *Butyricimonas* was enriched in NIH-Swiss **(Table S6).** These differences further suggest the strain-specific microbial associations with diet.

As an example of differential response, we compared how C57BI6/J and CD-1-associated microbiota differed. Based on beta-diversity analysis, these 2 strains had very similar microbial profiles, but C57BI6/J mice did not gain weight and showed no reduction in urinary ammonia excretion on the PD diet, while CD-1 mice both gained weight and showed a significant reduction in ammonia excretion on the PD diet, which suggests that CD-1 mice possibly are more efficient at utilizing the ammonia nitrogen for the production of amino acids. As these strains were apparently similar, yet had substantial differences in phenotypic response to a PD diet, we calculated which bacterial genera differentiated them. A non-parametric two-group test demonstrated that C57Bl6/J were enriched in *Oscillospira, Clostridium, and Coprococcus*, while CD-1 were significantly enriched in ESVs belonging to genus *Parabacteroides* and family Rickenellaceae (p_FDR_ < 0.05) (**Table S7**).

Random forest models were next employed to validate the bacterial signatures for the PD diet response identified using ANCOM. The random forest models built using the total data trained using strain type and diet type assignments, were able to predict the diet type (NC or PD) with an accuracy of 85% (OOB=0.15). Further, for diet type classification model, we identified ESVs that discriminate diet, such as *Akkermansia, Prevotella, Allobaculum, Turicibacter,* and *Parabacteroides*, which agrees with the non-parametric test results, validating the bacterial signatures that can be used to differentiate the strain type and diet type.

### Taxa-trait relationships

Next, we investigated how specific properties (weight, diet, or biochemical parameters) associated with taxa modules (i.e. group of correlated ESVs based on relative abundance). Using WGCNA, we identified five taxonomical modules which were arbitrarily assigned the colors yellow (11 ESVs), blue (13 ESVs), red (13 ESVs), turquoise (16 ESVs), and grey (34 ESVs) (see **Fig. 4A**). Using correlation cut-offs of 0.5 and p-value < 0.05, the blue module presented no significant association with any of the phenotypes tested. The yellow module was significantly positively correlated with strain type status and control or PD groups, and the taxonomic composition of this cluster agreed with the non-parametric ANCOM analysis. The red module, which included *Dehalobacterium, Anaerofustis, Roseburia, Oscillospira* and *Bilophila*, was positively correlated (p < 0.05) to the concentration of excreted urinary ammonia. Interestingly, the turquoise module including *Bacteroides, Parabacteroides, Mucispirillum, Enterococcus*, and *Allobaculum*, showed a significant but negative association with urinary ammonia (**Fig. 4A**). The grey module was significantly associated with strain status, group status and body weight. Creatinine concentration did not correlate with any of the modules.

**Figure 4:**
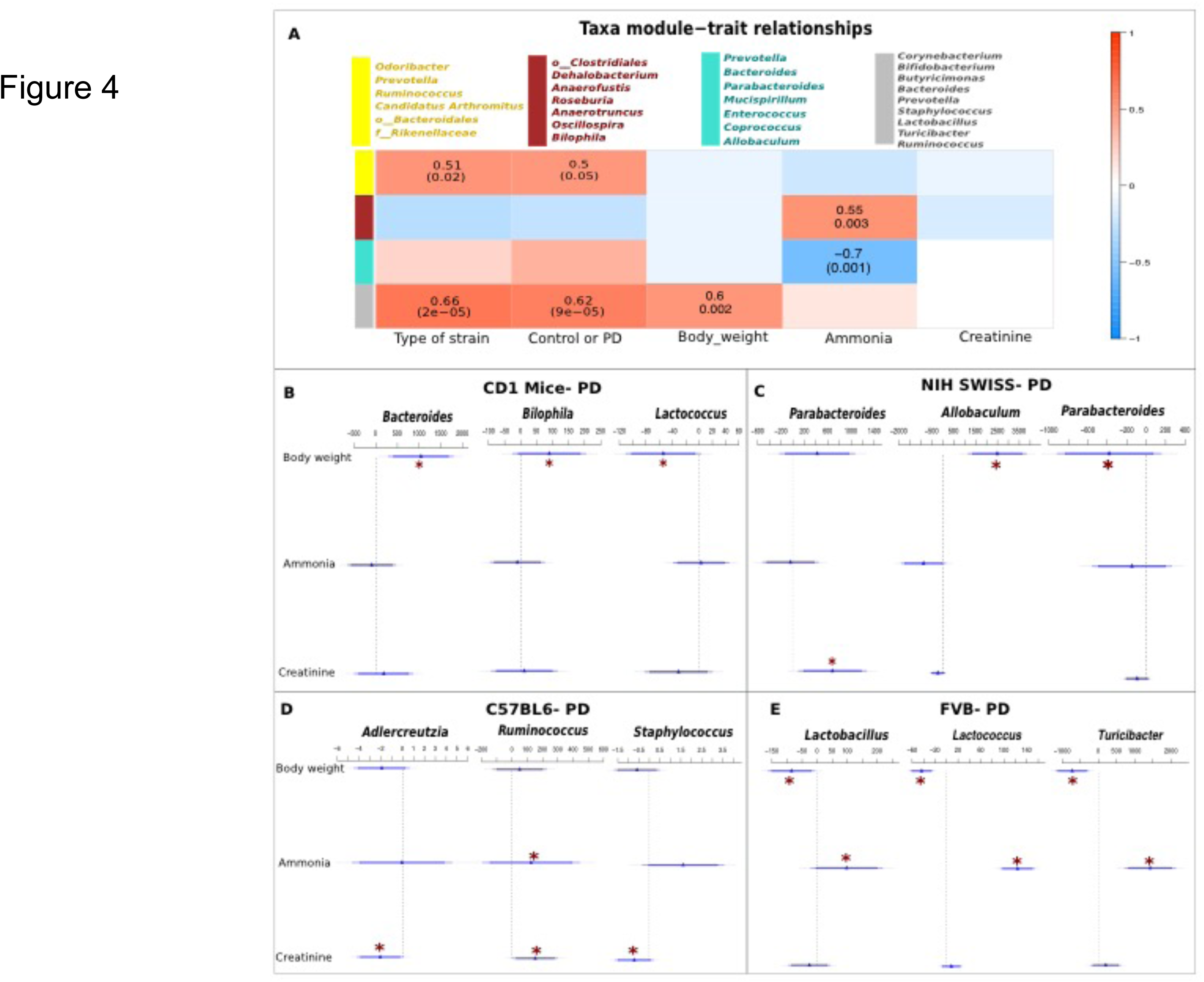
WGCNA analysis to assess the importance of module on a specific clinical trait. In the present figure, each row corresponds to a module eigengene, column to a trait. Each cell contains the corresponding correlation and p-value. The table is color coded by correlation according to the color legend (A). GLM model based on Poisson regression function was generated using the counts data for the differentially abundant genera for each PD-group (B-E), CD-1 (B), NIH-Swiss (C), C57BI6/J (D), and FVB (E). The significance of association between the genera and the body weight, ammonia and creatinine secretion was investigated using ANOVA (Chisq) test. The asterisks represent the significant correlations (positive or negative) with p value < 0.05.

GLM model fitting revealed significant associations between the PD diet microbiota in each of the 4 murine strains to body weight, urinary ammonia and creatinine excretion (**Fig. 4 B-E**). Across the CD-1 mice on PD, *Bacteroides* and *Bilophila* correlated positively (p < 0.05) with body weight, whereas, *Lactococcus* correlated negatively (p < 0.05) (**Fig. 4B**). *Parabacteroides* associated positively with creatinine (**Fig. 4B**). *Allobaculum* demonstrated positive correspondence with body weight, *Parabacteroides* showed negative correlation with body weight across the NIH-Swiss mice on PD (**Fig. 4C**). While for the C57BI6/J mice on PD, *Adlercreutzia* and *Staphylococcus* associated negatively (p < 0.05) with creatinine while *Ruminococcus* correlated positively with urinary ammonia as well as creatinine (**Fig. 4D**). Interestingly, across the FVB mice on PD, *Lactococcus, Lactobacillus* and *Turicibacter* correlated significantly with body weight (negative) and urinary ammonia (positive) (**Fig. 4E**).

### Functional association with protein deficient diet

We further assessed the predicted bacterial metabolic function of the microbiota and compared the predicted functional potential of the microbiota between individual strains on PD and NC diets (**Fig. 5**). Overall, the multi-group analyses between all the 4 strains demonstrated, a significant enrichment of predicted genes encoding the copper chaperon protein and cytochrome c-oxidase subunit II proteins in the FVB strain on the PD diet (**Fig. 5A, 5C**). The pyruvate dehydrogenase was significantly enriched in C57Bl6/J and NIH-Swiss strains on PD diet (**Fig. 5E**). Predicted genes encoding DNA excision repair protein ERCC-2 and sortase B were significantly enriched in the C57Bl6/J strain on PD diet (**Fig. 5D, 5F**). Two-group comparisons of the C57Bl6/J strain mice on NC and PD diets demonstrated significant enrichment of predicted genes encoding pyruvate dehydrogenase and thiamine biosynthase (**Fig. S1**). For the CD-1 strain, we identified predicted enzymes such as acetyl-CoA carboxylase, homoserine dehydrogenase, methyltransferases and ATP-dependent helicases to be significantly enriched on the PD diet (**Fig. S2**). For the FVB strain type, predicted enzymes such as thiamine biosynthesis, cytochrome oxidase and adenyltransferase were significantly enriched in the mice on PD diet (**Fig. S3**). Predicted genes encoding citrate lyase were significantly enriched in NIH-Swiss mice on the PD diet when compared to the NC diet (**Fig. S4**).

**Figure 5:**
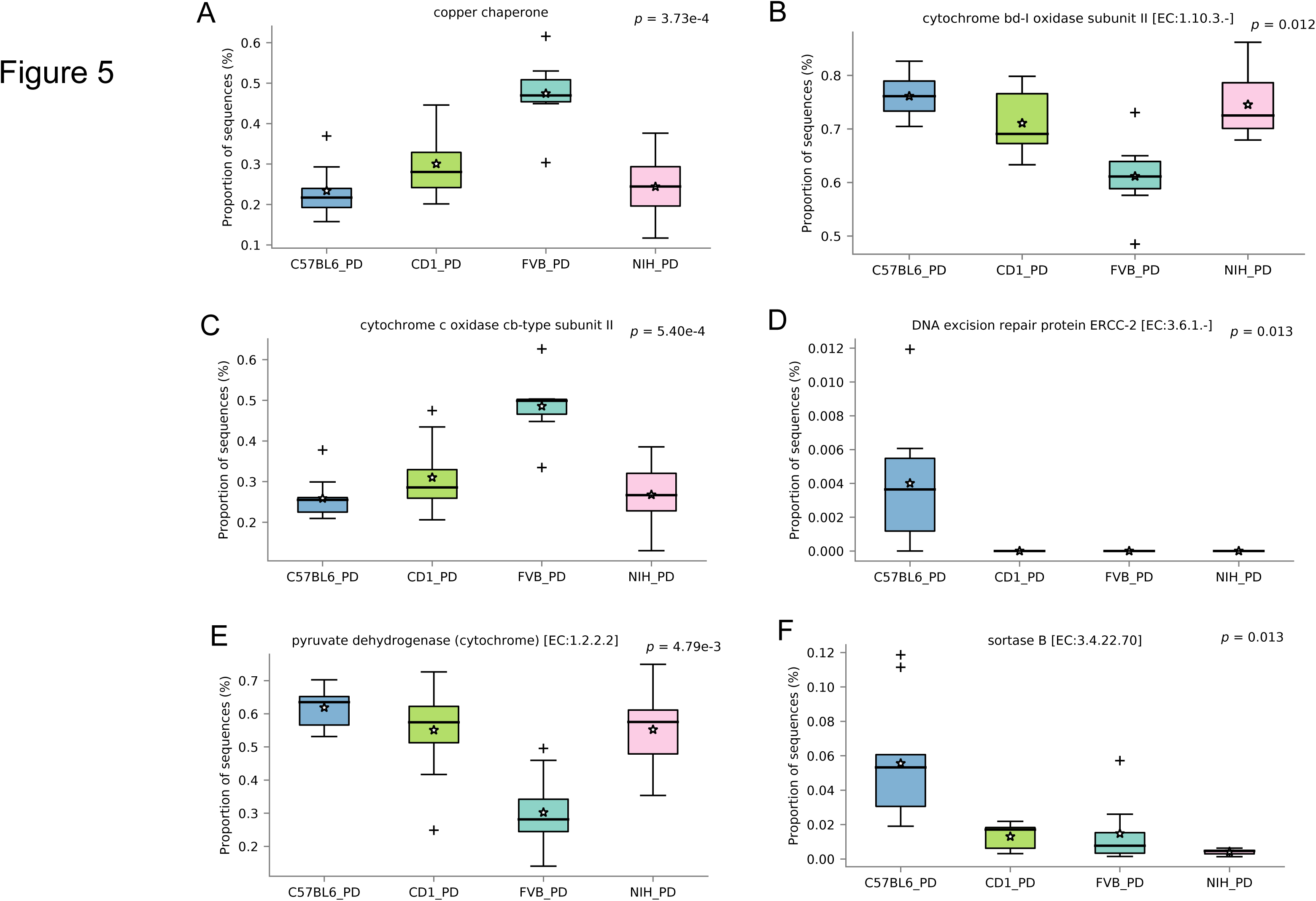
Significantly differential predicted bacterial functions (enzymes) between the 4 strains of mice i.e. CD-1, C57Bl6/J, FVB, NIH-Swiss on protein deficient diet (Fig A-F). All the genera were identified after applying Benjamini-Hochberg (BH) False Discovery Rate (FDR) correction to P-values (< 0.05).

Additionally, we compared the functional potential of CD-1 and C57Bl6/J strains on the PD diet to investigate the differential weight gain patterns of the 2 strains. The enzyme level annotations demonstrated significantly high abundance (p_FDR_ < 0.05) of bacterial dihydroorotate dehydrogenase, RNA-splicing ligase, and rRNA methyltransferases in CD-1 strains (**Fig. S5**). At the pathway level, an increased abundance for pathways involved in protein digestion/absorption and bile acid biosynthesis (primary and secondary) was observed in CD-1 strain compared to C57Bl6/J strains (**Fig. S6**).

## Discussion

Here we explored the hypothesis that different murine model strains (C57Bl6/J, CD-1, FVB, and NIH-Swiss) would have a different response to a protein restricted diet. Responses were quantified through both urinary N_2_ excretion and weight maintenance or growth. Microbial community structure and diversity were examined across murine strains and diet-associated responses. Overall, different microbial taxa were significantly correlated with responses to a restricted protein diet, which suggests that the microbiota may influence host-response during malnutrition events.

Growth presumes the existence of a metabolic state characterized by positive N_2_ balance, and thus serves as a summary index of the preservation of the capacity in an organism to sustain positive N_2_ balance on marginal protein intake. Here, we show that different gut microbial profiles correlated with growth on protein-restricted diets. C57BI6/J and NIH-Swiss mice on a 4 weeks PD diet failed to maintain the expected weight gain compared to their age matched controls on a NC diet. Interestingly, FVB and CD-1 mice retained comparable weekly weight gain between the PD and NC groups.

Protein breakdown provides more than 90% of amino acids to the metabolic pool and dietary protein intake provides much of the remainder. The main nitrogenous end products of catabolism of protein and amino acids are: urea, ammonia, and creatinine. Urinary excretion of these major moieties were measured as a marker of the status of N_2_ balance in the 4 strains of mice in response to a low protein feeding. The PD diet was associated with significantly diminished urinary and serum urea levels in CD-1 and C57BI6/J relative to NC fed mice, while in NIH-Swiss mice, urea was reduced in both urine and serum but only significantly in serum. FVB mice retained similar urea concentration both in urine and serum independent of diet. Significantly lower levels of urinary ammonia excretion in CD-1 mice and NIH-Swiss mice were induced by the PD feeding. However, in FVB mice, both NC and PD group demonstrated similar urinary ammonia excretion, while C57Bl6/J on the low protein diet exhibited only a slight reduction compared with NC fed animals. This observation suggests that the PD diet may have a mice strain-dependent impact in renal/urinary ammonia excretion. Serum ammonia levels, while lower on a PD diet, were not significantly reduced compared to NC diet for C57BI6/J, CD-1, and NIH-Swiss mice. It is interesting to note that serum ammonia concentration in the FVB mice was slightly greater on the PD diet compared with the NC diet. Renal creatinine excretion was quantified in urine at a single collection time point. The PD diet was associated with a consistent, but non-significant decrease in creatinine on a PD diet compared to an NC diet for all 4 strains of mice. Therefore, by quantifying urea, ammonia and creatinine, we have an initial analysis of the urinary N_2_ balance.

The possibility that bacteria can play a role in maintaining the N_2_ balance in the host during PD states has been previously suggested (23). The gut microbiota are responsible for producing nutrients for the host, including short chain fatty acids and essential amino acids (28). More recent evidence has demonstrated a specific correlative and causal association between microbial community structure, function and malnutrition. In 2013, it was shown that a fecal microbiota transplant into mice with samples from children suffering from Kwashiorkor resulted in weight loss and major nutrient metabolic changes in the mice (22). This same group has also begun to define how individual bacterial species from human gut microbiome respond to micronutrient deficient diet (8,26). A similar approach can be used to investigate PD diets, to help us define individual bacterial species of interest.

Gut microbes reside in the terminal ileum and cecum/ascending colon and utilize N_2_ principally in the form of ammonia (NH3) to synthesize amino acids for their own use (20). The origin of this NH3 nitrogen includes, dietary residue transiting the ileo-caecal valve to enter the caecum, gut secretions and sloughed gut mucosal cells; importantly, it includes a large flux of urea produced by the liver which enters the colon from the circulation (30). This N_2_ is used by the gut microbiota to trans-aminate non-essential amino acids and carbon skeletons as well as to fix the N_2_ in essential amino acids in the gut, which each can be passed into the circulation for use by the host. The magnitude of the utilization of urea N_2_ to make amino acids for the metabolic pool is determined by the interaction between demand and supply.

16S rRNA based gut microbiota analyses across the 4 strains of mice showed that alpha diversity was significantly reduced in mice on a PD diet compared to a NC diet. The multi-group ANCOM revealed 12 bacterial genera, including *Sutterella, Prevotella, Butyricimonas*, and *Enterococcus,* which significantly differentiated the four mice strains at baseline, i.e. independent of diet. Therefore, it is necessary to subtract the baseline differences when analyzing the PD-specific variations between the 4 strains. Each strain demonstrated a different set of genera that discriminated the mice on the PD diet compared to controls. For instance, PD diet led to significantly increased abundance of *Allobaculum* (C57Bl6/J), *Parabacteroides* (CD-1), *Turicibacter* (FVB) and *Mucispirillum* (NIH-Swiss). Further, WGCNA and GLM model fitting revealed significant association between the microbial variations and the body weight, ammonia and creatinine excretion. Genera such as *Bacteroides* and *Staphylococcus* showed a positive correlation with body weight, which has already been demonstrated (7). Similarly, genera such as *Bilophila, Roseburia, Oscillospira*, showed a significant association with the urinary ammonia excretion data.

The predicted functional potential of the bacterial communities showed significant differential abundance of predicted enzymes between the four strains. Interestingly, the PD diet increased the abundance of specific enzymes, including, ATP-dependent helicases, citrate lyase, and acetyl-CoA carboxylase when compared to the NC diet. Predicted genes encoding microbial pyruvate dehydrogenase increased significantly in C57Bl6/J mice on the PD diet when compared to NC diet, suggesting a physiological disruption leading to increased microbial cellular respiration. Previously, mice have shown an increase in muscle-based pyruvate dehydrogenase during significant loss of lean body mass when fasting (10,21). As such, it would appear that both host and microbial cells respond to protein restriction in a distinct way. Interestingly, the FVB mice, who retained weight on the PD diet, showed a significant reduction in the predicted abundance of microbial pyruvate dehydrogenase. Selective comparison of the CD-1 (gained weight) and C57Bl6/J (did not gain weight) strains on PD diet also demonstrated significant enrichment of dihydroorodate dehydrogenase, rRNA methyltransferases and RNA splicing ligase in the CD1 strains compared to C57Bl6/J strains.

This initial study presents some compelling evidence suggesting that differences in microbial community structure and prediction function may be associated with response to weight gain and N_2_ balance on a protein restricted diet. To go beyond this will require fecal microbiota transplant between strains to demonstrate phenotype transfer, then isolation of specific organisms associated with these outcomes and association in germ free animals to determine mechanistic interactions supporting outcomes. To our knowledge, this is the first study to report the association of murine microbial profiles with the ability to maintain positive N_2_ balance and during a protein deficient feeding.

## Acknowledgements

BTL is supported by the National Institutes of Health under award number, R01DK104927-01A1 and Department of Veterans Affairs, Veterans Health Administration, Office of Research and Development, Career Development (Grant no. 1IK2BX001587-01).

## References

1. Amir, Amnon, Daniel McDonald, Jose A. Navas-Molina, Evguenia Kopylova, James T. Morton, Zhenjiang Zech Xu, Eric P. Kightley, et al. 2017. “Deblur Rapidly Resolves Single-Nucleotide Community Sequence Patterns.” MSystems 2 (2). https://doi.org/10.1128/mSystems.00191-16.

2. Badaloo, Asha, Marvin Reid, Terrence Forrester, William C. Heird, and Farook Jahoor. 2002. “Cysteine Supplementation Improves the Erythrocyte Glutathione Synthesis Rate in Children with Severe Edematous Malnutrition.” The American Journal of Clinical Nutrition 76 (3): 646–52. https://doi.org/10.1093/ajcn/76.3.646.

3. Black, Robert E., Saul S. Morris, and Jennifer Bryce. 2003. “Where and Why Are 10 Million Children Dying Every Year?” The Lancet 361 (9376): 2226–34. https://doi.org/10.1016/S0140-6736(03)13779-8.

4. Bokulich, Nicholas A., Sathish Subramanian, Jeremiah J. Faith, Dirk Gevers, Jeffrey I. Gordon, Rob Knight, David A. Mills, and J. Gregory Caporaso. 2013. “Quality-Filtering Vastly Improves Diversity Estimates from Illumina Amplicon Sequencing.” Nature Methods 10 (1): 57–59. https://doi.org/10.1038/nmeth.2276.

5. Caporaso, J Gregory, Justin Kuczynski, Jesse Stombaugh, Kyle Bittinger, Frederic D Bushman, Elizabeth K Costello, Noah Fierer, et al. 2010. “QIIME Allows Analysis of High-Throughput Community Sequencing Data.” Nature Methods 7 (5): 335–36. https://doi.org/10.1038/nmeth.f.303.

6. Chiu, Chun-Huo, and Anne Chao. 2016. “Estimating and Comparing Microbial Diversity in the Presence of Sequencing Errors.” PeerJ 4: e1634. https://doi.org/10.7717/peerj.1634.

7. Clarke, Siobhan F., Eileen F. Murphy, Kanishka Nilaweera, Paul R. Ross, Fergus Shanahan, Paul W. O’Toole, and Paul D. Cotter. 2012. “The Gut Microbiota and Its Relationship to Diet and Obesity.” Gut Microbes 3 (3): 186–202. https://doi.org/10.4161/gmic.20168.

8. Crawford, Peter A., Jan R. Crowley, Nandakumar Sambandam, Brian D. Muegge, Elizabeth K. Costello, Micah Hamady, Rob Knight, and Jeffrey I. Gordon. 2009. “Regulation of Myocardial Ketone Body Metabolism by the Gut Microbiota during Nutrient Deprivation.” Proceedings of the National Academy of Sciences of the United States of America 106 (27): 11276–81. https://doi.org/10.1073/pnas.0902366106.

9. DeBoer, Mark D., Aldo A. M. Lima, Reinaldo B. Oría, Rebecca J. Scharf, Sean R. Moore, Max A. Luna, and Richard L. Guerrant. 2012. “Early Childhood Growth Failure and the Developmental Origins of Adult Disease: Do Enteric Infections and Malnutrition Increase Risk for the Metabolic Syndrome?” Nutrition Reviews 70 (11): 642–53. https://doi.org/10.1111/j.1753-4887.2012.00543.x.

10. Fields, A. L., N. Falk, S. Cheema-Dhadli, and M. L. Halperin. 1987. “Accelerated Loss of Lean Body Mass in Fasting Rats Due to Activation of Pyruvate Dehydrogenase by Dichloroacetate.” Metabolism: Clinical and Experimental 36 (7): 621–24.

11. Flores, Gilberto E., Scott T. Bates, Dan Knights, Christian L. Lauber, Jesse Stombaugh, Rob Knight, and Noah Fierer. 2011. “Microbial Biogeography of Public Restroom Surfaces.” PLoS ONE 6 (11). https://doi.org/10.1371/journal.pone.0028132.

12. Galler, Janina R., Cyralene P. Bryce, Miriam L. Zichlin, Garrett Fitzmaurice, G. David Eaglesfield, and Deborah P. Waber. 2012. “Infant Malnutrition Is Associated with Persisting Attention Deficits in Middle Adulthood.” The Journal of Nutrition 142 (4): 788–94. https://doi.org/10.3945/jn.111.145441.

13. Galler, Janina R., Cyralene P. Bryce, Miriam L. Zichlin, Deborah P. Waber, Natalie Exner, Garrett M. Fitzmaurice, and Paul T. Costa. 2013. “Malnutrition in the First Year of Life and Personality at Age 40.” Journal of Child Psychology and Psychiatry, and Allied Disciplines 54 (8): 911–19. https://doi.org/10.1111/jcpp.12066.

14. Gelman, Andrew. 2008. “Scaling Regression Inputs by Dividing by Two Standard Deviations.” Statistics in Medicine 27 (15): 2865–73. https://doi.org/10.1002/sim.3107.

15. Horvath, Steve, and Jun Dong. 2008. “Geometric Interpretation of Gene Coexpression Network Analysis.” PLoS Computational Biology 4 (8). https://doi.org/10.1371/journal.pcbi.1000117.

16. Langfelder, Peter, and Steve Horvath. 2008. “WGCNA: An R Package for Weighted Correlation Network Analysis.” BMC Bioinformatics 9 (December): 559. https://doi.org/10.1186/1471-2105-9-559.

17. Lozupone, Catherine, Manuel E Lladser, Dan Knights, Jesse Stombaugh, and Rob Knight. 2011. “UniFrac: An Effective Distance Metric for Microbial Community Comparison.” The ISME Journal 5 (2): 169–72. https://doi.org/10.1038/ismej.2010.133.

18. Mandal, Siddhartha, Will Van Treuren, Richard A. White, Merete Eggesbø, Rob Knight, and Shyamal D. Peddada. 2015. “Analysis of Composition of Microbiomes: A Novel Method for Studying Microbial Composition.” Microbial Ecology in Health and Disease 26 (May). https://doi.org/10.3402/mehd.v26.27663.

19. McMurdie, Paul J., and Susan Holmes. 2013. “Phyloseq: An R Package for Reproducible Interactive Analysis and Graphics of Microbiome Census Data.” PloS One 8 (4): e61217. https://doi.org/10.1371/journal.pone.0061217.

20. Metges, Cornelia C. 2000. “Contribution of Microbial Amino Acids to Amino Acid Homeostasis of the Host.” The Journal of Nutrition 130 (7): 1857S–1864S. https://doi.org/10.1093/jn/130.7.1857S.

21. Rinnankoski-Tuikka, Rita, Mika Silvennoinen, Sira Torvinen, Juha J. Hulmi, Maarit Lehti, Riikka Kivelä, Hilkka Reunanen, and Heikki Kainulainen. 2012. “Effects of High-Fat Diet and Physical Activity on Pyruvate Dehydrogenase Kinase-4 in Mouse Skeletal Muscle.” Nutrition & Metabolism 9 (June): 53. https://doi.org/10.1186/1743-7075-9-53.

22. Smith, Michelle I., Tanya Yatsunenko, Mark J. Manary, Indi Trehan, Rajhab Mkakosya, Jiye Cheng, Andrew L. Kau, et al. 2013. “Gut Microbiomes of Malawian Twin Pairs Discordant for Kwashiorkor.” Science (New York, N.Y.) 339 (6119): 548–54. https://doi.org/10.1126/science.1229000.

23. Smythe, P. M. 1958. “CHANGES IN INTESTINAL BACTERIAL FLORA AND ROLE OF INFECTION IN KWASHIORKOR.” The Lancet 272 (7049): 724–27. https://doi.org/10.1016/S0140-6736(58)91336-9.

24. Tennant, Ingrid A., Alan T. Barnett, Debbie S. Thompson, Jan Kips, Michael S. Boyne, Edward E. Chung, Andrene P. Chung, et al. 2014. “Impaired Cardiovascular Structure and Function in Adult Survivors of Severe Acute Malnutrition.” Hypertension (Dallas, Tex.: 1979) 64 (3): 664–71. https://doi.org/10.1161/HYPERTENSIONAHA.114.03230.

25. Thompson, Luke R., Jon G. Sanders, Daniel McDonald, Amnon Amir, Joshua Ladau, Kenneth J. Locey, Robert J. Prill, et al. 2017. “A Communal Catalogue Reveals Earth’s Multiscale Microbial Diversity.” Nature 551 (7681): 457–63. https://doi.org/10.1038/nature24621.

26. Turnbaugh, Peter J., Vanessa K. Ridaura, Jeremiah J. Faith, Federico E. Rey, Rob Knight, and Jeffrey I. Gordon. 2009. “The Effect of Diet on the Human Gut Microbiome: A Metagenomic Analysis in Humanized Gnotobiotic Mice.” Science Translational Medicine 1 (6): 6ra14. https://doi.org/10.1126/scitranslmed.3000322.

27. Victora, Cesar G., Linda Adair, Caroline Fall, Pedro C. Hallal, Reynaldo Martorell, Linda Richter, Harshpal Singh Sachdev, and Maternal and Child Under nutrition Study Group. 2008. “Maternal and Child Undernutrition: Consequences for Adult Health and Human Capital.” Lancet (London, England) 371 (9609): 340–57. https://doi.org/10.1016/S0140-6736(07)61692-4.

28. Wong, Julia M. W., Russell de Souza, Cyril W. C. Kendall, Azadeh Emam, and David J. A. Jenkins. 2006a. “Colonic Health: Fermentation and Short Chain Fatty Acids.” Journal of Clinical Gastroenterology 40 (3): 235.

29. Wong, Julia M. W., Russell de Souza, Cyril W. C. Kendall, Azadeh Emam, and David J. A. Jenkins. 2006b. “Colonic Health: Fermentation and Short Chain Fatty Acids:” Journal of Clinical Gastroenterology 40 (3): 235–43. https://doi.org/10.1097/00004836-200603000-00015.

30. Yang, Yu-Xiang, Zhao-Lai Dai, and Wei-Yun Zhu. 2014. “Important Impacts of Intestinal Bacteria on Utilization of Dietary Amino Acids in Pigs.” Amino Acids 46 (11): 2489–2501. https://doi.org/10.1007/s00726-014-1807-y.

